# Shape-constrained, changepoint additive models for time series omics data with cpam

**DOI:** 10.1101/2024.12.22.630003

**Authors:** Luke A. Yates, Jazmine L. Humphreys, Michael A. Charleston, Steven M. Smith

## Abstract

Time series omics experiments are critical for the study of a wide range of biological processes such as cell differentiation and developmental programs or responses to pathogens and environmental cues. While statistical tools for differential analysis across static conditions have matured, comparable comprehensive methodology is lacking for time series data. Here, we introduce `cpam`, a novel time series method and user-friendly software that performs temporal differential analysis of omics data for case-only or case-control time series, incorporating quantification uncertainty. Powerful features of `cpam` include changepoint detection and shape-constrained temporal trend estimation to allocate omics data into similar clusters. Performance evaluation shows that `cpam` outperforms existing time series methods in terms of control of the false discovery rate versus power to detect temporal changes and accurate changepoint estimation. In particular, we find that trend-based models provide superior performance compared to pairwise comparisons, even with as few as six time points. The software provides an interactive interface with modern customizable visualisations, offering both graphical and statistical insight into complex molecular processes. Application to published transcriptome data illustrates RNA isoform-level modelling together with high resolution clustering during human embryogenesis and the identification of 910 novel genes differentially expressed in response to excess light in Arabidopsis. This method has the potential for application to a range of omics data, including single-cell analyses via pseudobulked data, providing the analytical power to reveal complex temporal patterns of gene expression in any biological system.

## 1 Introduction

Analysing temporal changes in genome-wide gene expression is fundamentally important for the study of many biological processes, such as the progression of developmental programs or responses to external factors such as nutrients, xenobiotics, pathogens and environmental signals (Snead and Clark, 2022). Omics technologies for the analysis of chromatin, transcriptome, proteome and metabolome have developed rapidly such that they are increasingly powerful and affordable, generating large amounts of data in single experiments (Vandereyken et al., 2023). Thus, a major limiting factor now lies in our ability to rapidly and effectively analyse such temporal data to visualise patterns of gene expression and to deduce underlying pathways and networks.

Methods for analysing time series omics data can be categorised based on the type of temporal trend being inferred. In the simplest case, trends are eschewed altogether, and each time point is treated independently using pairwise comparisons or treating time as a categorical factor. Despite the simplicity of the pairwise approach, it can be effective when there are few time points or when observations are sufficiently separated in time to be effectively independent (Spies et al., 2019).

The notion of temporal autocorrelation—the extent to which measurements that are recorded closely in time tend to be similar in value—is fundamental to the analysis of time series data. This concept is important because temporally autocorrelated data are not independent, and this dependence must be taken into account to ensure valid statistical inference. In the context of genomic networks, temporal autocorrelation is expected and various approaches have been used to model it. These methods range from linear and polynomial trends implemented via commonly used software such as DESeq2 (Love et al., 2014) and edgeR (McCarthy et al., 2012) to more complex approaches such as (linear) changepoint models (Bacher et al., 2018), combinations of fixed-form trends (Fischer et al., 2018; Bar et al., 2022), and non-parametric techniques such as Gaussian processes and penalised splines that force trends to change smoothly over time (Nueda et al., 2014; Straube et al., 2015; Topa and Honkela, 2018).

In a broader context beyond time series, best practices in statistical genomics have continuously evolved over the past 20 years. The high-throughput revolution has produced a plethora of ‘small *n* large *p*’ data where the number of samples (*n*) is low while the number of statistical comparisons (*p*) is extremely high (Rice et al., 2008). This situation exacerbates the perennial challenge of discovery and replicability in the sciences, which rely on rigorous statistical methods to control type I error rates while maintaining sufficient power to detect true effects (Benjamini and Hochberg, 1995). Variance estimation, which is critical for valid inference, is particularly challenging given the typical sample sizes of genomics data. Modern regularisation (also called shrinkage or moderation) techniques that allow the sharing of information across genes, have emerged as an effective means to mitigate this issue (Robinson and Smyth, 2007; Van De Wiel et al., 2012; Love et al., 2014).

A further challenge is that gene products can occur as different variants, particularly apparent in the case of transcripts and proteins, usually referred to as RNA and protein isoforms. In many cases in complex eukaryotes, a single gene can produce multiple RNA isoforms, potentially encoding protein isoforms. In the case of RNA isoforms, quantification is often uncertain, because of mapping ambiguities between (short) reads and the available reference transcriptome (Soneson et al., 2015). Recently developed methods estimate this quantification uncertainty (Bray et al., 2016; Patro et al., 2017) and allow it to propagate into subsequent statistical analysis, improving inference (Pimentel et al., 2017; Zhu et al., 2019; Baldoni et al., 2023). A related consideration is that statistical analysis at the transcript level, followed by the aggregation of the *p* values from the transcript to the gene level, has more power to discover differentially active genes than if the data themselves are aggregated to the gene level prior to analysis (Yi et al., 2018). The former can detect important differences such as differential RNA isoform abundance (e.g., between experimental conditions or across a time series), which may be obscured if the data are aggregated.

Current time series omics methods struggle to accurately infer biological signals that can involve both gradual and sharp changes in abundance of gene products while maintaining statistical rigour. Pairwise comparisons treat timepoints as independent, ignoring temporal structure, which significantly reduces statistical power. Linear trends, even with changepoints, and other parametric models tend to impose overly simplistic functional forms that cannot capture complex regulatory dynamics (Tonner et al., 2016). Existing non-parametric approaches use smoothing constraints to improve on polynomial regression for nonlinear trends, but are unable to accommodate step changes in gene expression. Most time series methods do not incorporate quantification uncertainty or share information between genes.

To address these challenges, we present `cpam`, an R package for omics time series analysis that combines current best practices in statistical genomics with biologically motivated innovations. `cpam` introduces changepoint additive models that can detect sharp transitions using changepoints, while also modelling smooth expression changes. The method includes shape-constrained trends that cluster genes or transcripts into biologically meaningful temporal shape classes. Here, we focus on its application to transcriptome data, but recognise that it can equally be adapted for other omics platforms. `cpam` performs gene-or transcript-level inferences while accounting for quantification uncertainty, aggregating *p*-values at the gene level, and sharing information across genes when estimating dispersions. In comprehensive benchmarking, `cpam` demonstrates superior performance through well-calibrated *p*-values, improved power to detect true positives while controlling false detection, and accurate changepoint detection compared to existing methods. `cpam` provides clear and interactive visualisations of temporal dynamics and changes in gene and transcript abundance, essential for the interpretation of complex genome-wide time series experiments.

`cpam` is freely available from the repository at https://github.com/l-a-yates/cpam

## 2 Results

### Statistical model and inference

The `cpam` model sets out to analyse time series omics data in three stages so that the later computationally intensive stages focus on a subset of differentially expressed genes. Stage 1 identifies differentially expressed genes through analysis of their products, which for current considerations is provided by transcriptome data. In this case, the RNA products can be aggregated at the gene level or considered at the level of individual RNA isoforms. Before modelling, data are filtered to exclude zero-count and low-count genes (see Methods for details). Stage 2 involves the determination of change points for the differentially expressed RNAs. Stage 3 involves selecting temporal shapes to which each RNA conforms to facilitate subsequent clustering analysis. In addition, unconstrained shapes can be visualised and smoothed as required. Hereafter we refer to isoforms (or genes for a gene-level analysis) as the ‘*targets*’ of computational analysis.

### Stage 1: Temporal differential expression of genes and isoforms

In the first modelling stage, we fit penalised smoothing splines with a negative binomial response distribution. A spline is a smooth curve that interpolates the observed data (see Figure 1A for examples). For observed counts *y*_*ij*_, indexed by target *i* and sample *j*, the Stage 1 statistical model is

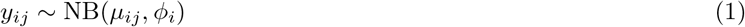

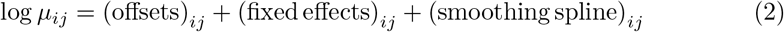

where *µ*_*ij*_ is the modelled mean, *ϕ*_*i*_ is the dispersion parameter, and NB is the negative binomial probability mass function. The optional term (offsets)_*ij*_ includes, by default, weights to account for unequal sampling depths and a possible scaling factor to account for uncertainty in the quantification of transcripts (Baldoni et al., 2023) (see Figure 2A). The term (fixed effects)_*ij*_ includes an intercept and an optional treatment effect for case-control series, in addition to any user-provided covariates, such as batch identifiers. The term (smoothing spline)_*ij*_ comprises one (for case-only) or two (for case-control) thin-plate temporal smoothing splines each with an estimated smoothing parameter, which controls trend smoothness (see Figure 1B). The dispersion estimates are regularised using an empirical Bayes approach (McCarthy et al., 2012). For further details, see *Methods*.

**Fig 1.**
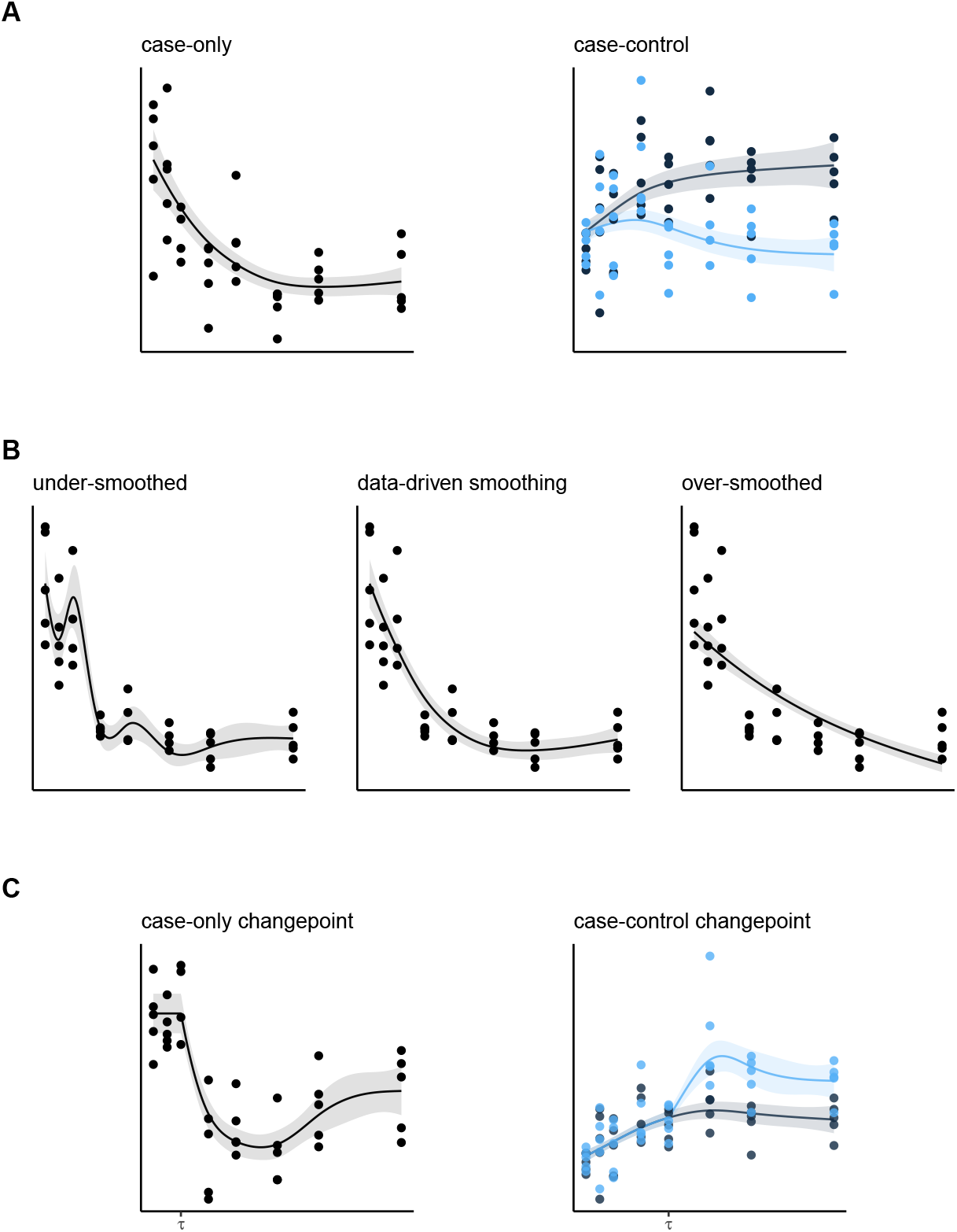
Spline-based models used in cpam. **A** Example smoothing splines. **B** Effect of the smoothing parameter on spline curvature. The middle plot shows the data-driven method used in `cpam`. **C** Example splines with a changepoint. For all plots, the x-axes denote time and the y-axes the response. The points are observed data and the solid lines and filled envelopes are the estimated smoothing splines and their standard errors, respectively. The blue features denote a treatment group.

**Fig 2.**
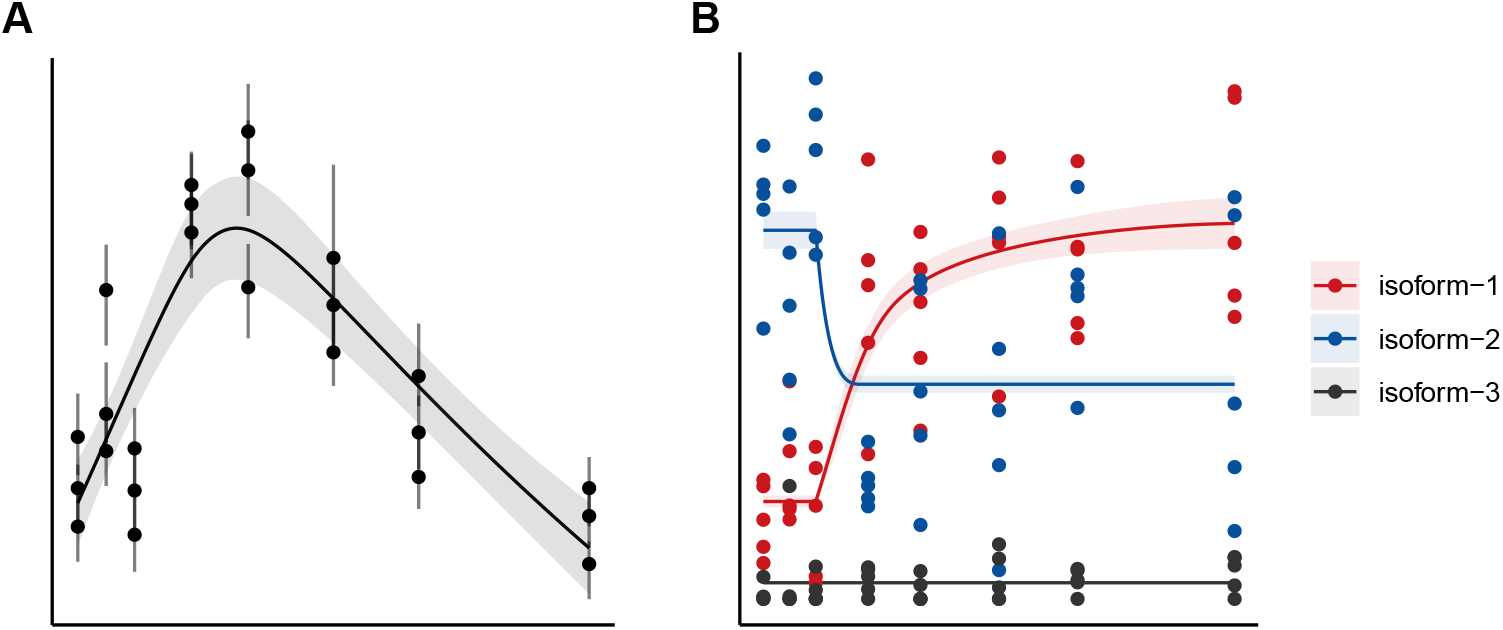
Data and trends for genes with multiple RNA isoforms.**A** Isoform quantification uncertainty. Mapping ambiguities generate uncertainty in the estimated number of counts at the isoform level. When estimated (e.g., using bootstrapping), `cpam` can plot these uncertainties and takes them into account when fitting models and making inferences. **B** Visualising trends for multiple gene isoforms. `cpam` detects and plots temporal processes such as isoform switching and differential transcript usage. The points and vertical lines are the mean and the 63.8% bootstrapped interval of the observed data, respectively. The solid lines and envelopes are the estimated smoothing splines and their standard errors.

The statistical significance of the smoothing spline (1) for each target is assessed by comparison with a null model. In case-only series, the null model has a constant mean over time (i.e., a horizontal line), while in case-control series, the null model uses a single shared spline for both case and control, rather than the alternative of a separate spline for each (see Figure 1A). Although accurately estimating *p*-values for penalised additive models is challenging (Wood, 2017), using a negative binomial distribution instead of a Gaussian model of logged counts, along with a separate estimate of the smoothing parameter for each target, resulted in well-calibrated *p*-values for the simulated time series data (see *Simulation studies for performance benchmarking* for calibration results). This approach uses a generalised likelihood ratio test that takes into account the smoothing procedure and the effective complexity of the model (Wood, 2012).

Modelling omics data at the RNA isoform level greatly increases the number of targets compared to gene-level counts, leading to more statistical comparisons and reduced power with classical *p*-value adjustments. To address this, `cpam` employs *p*-value aggregation, combining *p*-values from isoforms into a single *p*-value for each gene, which increases statistical power over gene-aggregated models while still allowing inference of differential transcript levels (Lancaster, 1961; Yi et al., 2018). The gene-level *p*-values are then adjusted for multiple comparisons to assess differential expression at gene and RNA isoform levels over time and treatments. The default adjustment in `cpam` is the false discovery rate (FDR), or Benjamini-Hochberg (BH) method (Benjamini and Hochberg, 1995), with other options available. A subset of differentially expressed genes is identified by choosing a threshold *α* for discovery; e.g., *p*_adjusted_ ≤ *α*.

### Stage 2: Changepoint detection

A changepoint is a point in a time series where there is an abrupt change in the statistical behaviour of the data, such as a change from constant to time-varying counts or a change in variability. Although gene networks typically exhibit smooth changes (e.g., due to self-regulating feedback), rapid changes can occur due to threshold effects (e.g., attainment of a critical concentration of a transcription factor) or switch-like dynamics (e.g., due to positive feedback), particularly in response to new conditions.

We extend the spline models of the previous section to include an initial period of constant (case-only) or common (case-control) mean counts followed by smooth timevarying counts. Example trajectories are shown in Figure 1C. The changepoint model augments (2) and is written as

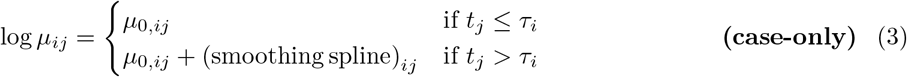

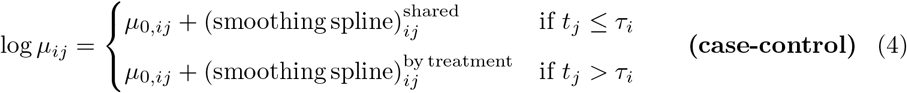

where *µ*_0,*ij*_ = (offsets)_*ij*_ +(fixed effects)_*ij*_ , *t*_*j*_ is the time point of sample *j*, and *τ*_*i*_ is the changepoint for target *i*. We limit *τ*_*i*_ to the discrete observed values in the series, since, despite the use of smoothing constraints, there is generally insufficient information to infer the timing of changepoints beyond the temporal resolution of the data. Although we generally do not anticipate the true temporal trends to involve such abrupt changes in slope, changepoint models are a good approximation to rapidly changing continuous processes given the limitation of discrete sampling.

To estimate changepoints, we fit models (3) or (4) for each value of *τ*_*i*_ and select among them using generalised cross-validation (GCV, default) or the Akaike Information Criterion (AIC). Model-selection uncertainty can be visualised by plotting the mean and standard errors of pointwise score differences with respect to the best-scoring model (see Figure 10). These standard errors can be used to apply the one-standard-error rule, to avoid overfitting by selecting the least complex model (i.e., the largest *τ*_*i*_) within one standard error of the best model (Yates et al., 2021). See *Methods* for details.

Given the computational cost of fitting a separate model for each candidate changepoint, `cpam` only estimates changepoints for targets associated with ‘significant’ genes at the chosen threshold *α*. In the limiting case *α* = 1, a changepoint is estimated for every target. These estimated changepoints provide a natural grouping variable, aiding downstream analyses such as clustering, gene-network inference (Liang and Kelemen, 2018; Marku and Pancaldi, 2023), or Granger causality (Deshpande et al., 2022).

### Stage 3: Shape selection

The final modelling stage involves selecting shape-constrained trends to cluster targets based on their temporal trajectories. These shapes expand upon traditional up- and down-regulated classes by capturing trends such as rising then falling, or rising then stabilising. To mathematically characterise candidate shapes, we use shape-constrained additive models (Pya and Wood, 2014) which extend the class of changepoint additive models (3) and (4). Shape-constrained splines can be convex, concave, or unconstrained, and may be monotonically increasing, decreasing, or neither. For targets with a changepoint near the series end or a linear-time response, candidate shapes are increasing or decreasing linear. Figure 3 displays examples of shape-constrained trends plotted on the log scale. Model estimation on the log scale means that shape selection and subsequent clustering is easily decoupled from the magnitude of the response.

**Fig 3.**
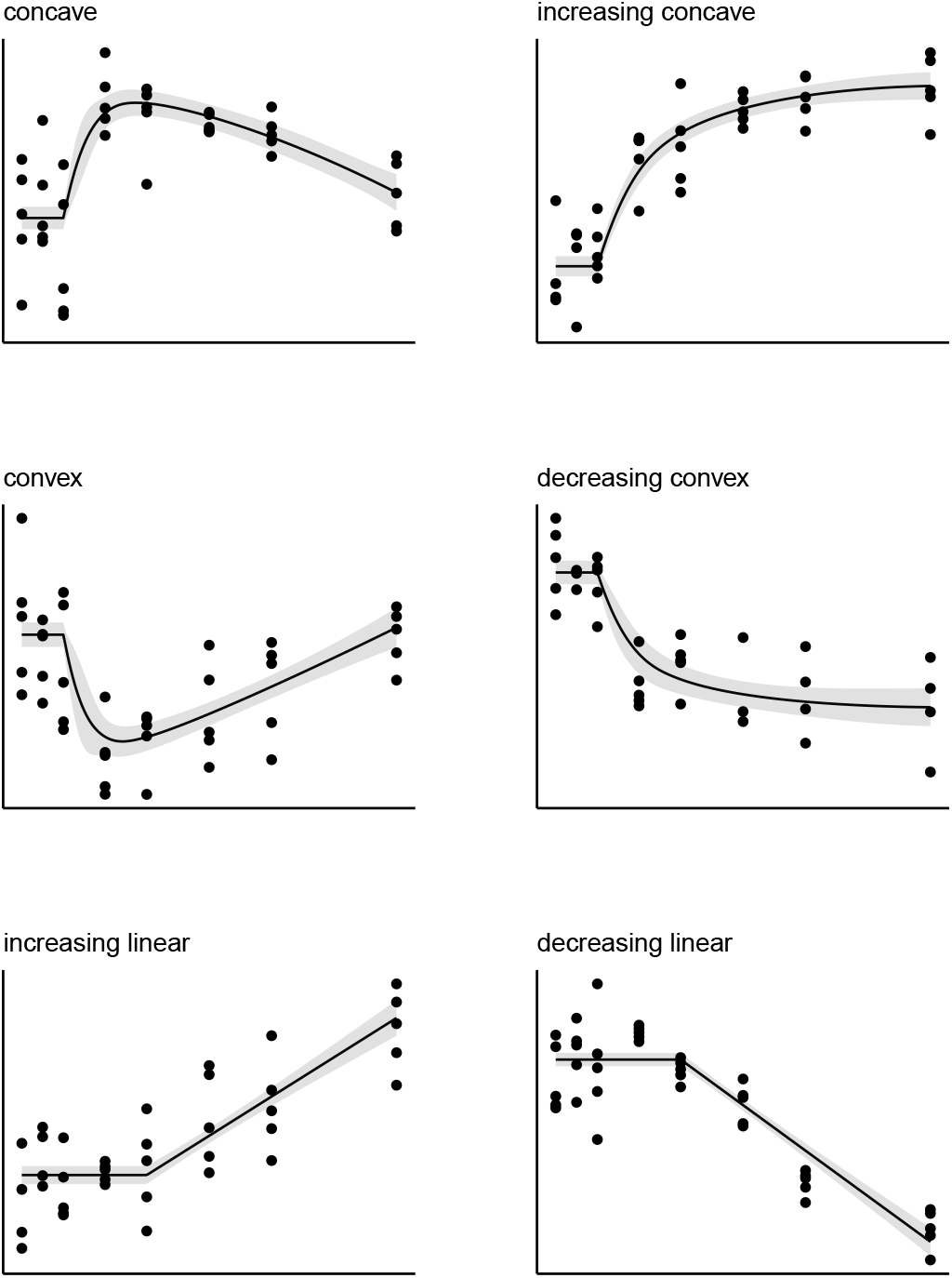
Prototypical temporal expression patterns modelled by `cpam`. Each panel shows a different shape-constrained pattern that can occur after a changepoint: concave decreasing/increasing (top row), convex increasing/decreasing (middle row), and linear increasing/decreasing (bottom row).

We chose this class of shapes for their flexibility and ability to represent biologically significant trends. Instead of relying on fixed parametric forms such as exponentially decaying or sigmoidal functions, which may poorly approximate underlying trajectories (Tonner et al., 2016), shape-constrained splines provide interpretable shapes by restricting only the sign of the first and second derivatives. These shapes were chosen to represent diverse biological dynamics for a wide range of biological processes.

For example, transient up- and down-regulation (concave and convex shapes respectively), transition to a different steady state under new conditions (monotonic convex or concave) or sustained up- or down-regulation throughout the time series (linear).

The changepoint estimated by `cpam` is an important component of the overall temporal shape. For instance, a constant trend changing to an increasing concave resembles a sigmoid, but estimates a specific time for the initial change and allows for more flexibility in the latter trajectory. The estimated changepoints can be used to cluster targets within a shape-constrained class. Figure 4 illustrates clusters based on changepoint and shape characteristics using `cpam`.

**Fig 4.**
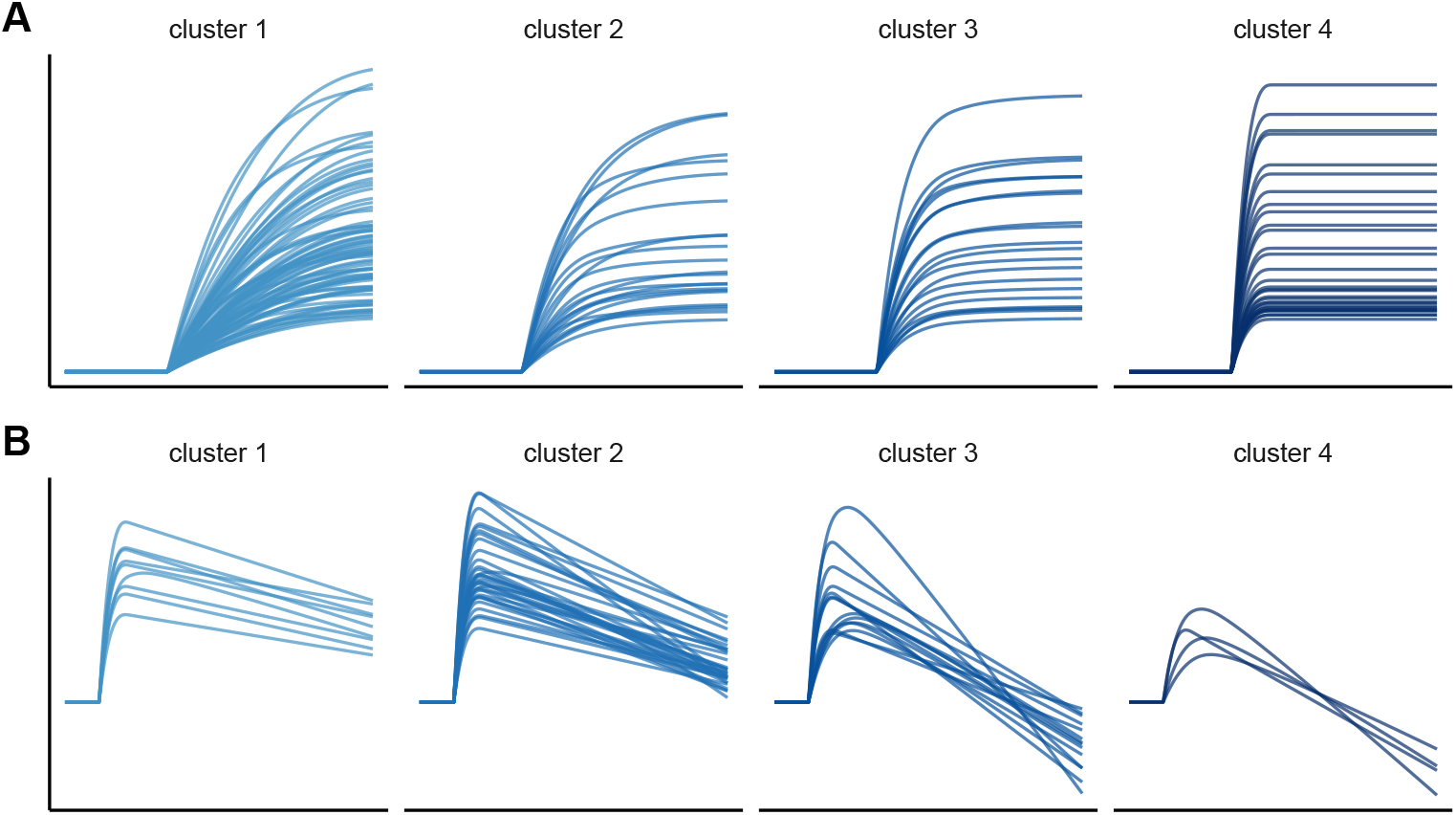
Examples of shape-based clustering using `cpam`.**A** Targets filtered by changepoint and shape (all increasing concave), and clustered by increasing stabilisation rate (time to reach 90% of the maximum response). **B** Targets filtered by changepoint, shape (concave) and time of maximum response (all the same), and clustered by the extent of down-regulation post maxima (ratio of the maximum to the final response). All trends are plotted on a logged vertical scale

Using smoothed temporal trends, shape-constrained additive models de-noise the data and cluster targets into distinct, interpretable shapes (Bar et al., 2022). In comparison, clustering unmodelled data (e.g., *k*-means) often generates hard-to-interpret groups of erratic trajectories (Humphreys et al., 2024). Even algorithmic clustering of modelled outputs such as polynomial and smoothing spline regression coefficients often produces clusters that mix divergent trend types.

Two shape selections for each target are made by `cpam`, the first selecting among linear, convex and concave shape classes and their monotonic variants, and the second selecting among the first options plus an ‘unconstrained’ smooth. The inclusion of the ‘unconstrained’ type provides the flexibility to detect targets beyond simpler trends. Shapes are selected using either generalised cross validation (GCV, default) or the Akaike Information Criterion (AIC). For computational reasons, as per the changepoint estimation, `cpam` only selects shapes for those genes, or their isoforms, identified as significant at the chosen FDR threshold *α*.

### Simulation studies for performance benchmarking

We evaluated `cpam` against several alternative methods under diverse simulation conditions and goals (Brooks et al., 2024). Alternatives were selected for their past benchmark performance, relevance to research goals, widespread genomic use, and software availability. We included time-as-a-factor (factor) and static pairwise comparisons (pairwise) as additional methods fitted using DESeq2 (Love et al., 2014). For the latter, the *p*-value for each target was set to the lowest pairwise *p*-value, thus marking targets as significant if any adjusted *p*-value was equal to or below the nominal threshold at any timepoint. FDR correction was applied to the aggregated set of all *p*-values.

We simulated omics count data from a human RNA-Seq dataset (Cheung et al., 2010) following the approach used in the comparative study of time series methods by Spies et al. (2019). The processed count data were obtained form the Recount resource (Frazee et al., 2011). Briefly, we randomly sampled 30,000 mean-dispersion pairs from an empirical distribution model. Replicates from a pair-specific NB distribution were simulated, averaging *n* = min(500, mean*/*4) samples to minimise the influence of extreme values, generating biologically plausible data. Temporal patterns were created by multiplying simulated replicates by predicted values from shape-constrained additive models for randomly selected shapes. This method simulated case-only and case-control data sets for short (6 time points) and long (12 timepoints) series. Additional simulation parameters included the number of replicates (baseline = 3) and the maximum expected log-base-2 fold change (LFC) with respect to the first time point for non-null targets, set to low (LFC = 1, baseline) or high (LFC = 2).

Simulating data from empirically informed shape-constrained additive models produces biologically plausible time series trajectories. However, this shape-based approach does not favour `cpam` over alternative methods when benchmarking *p*-value distributions, FDR control, and target discovery, since *p*-value calculations and significance assessments in `cpam` occur before the use of shape-based modelling.

### Distribution of *p*-values under the null hypothesis

Calculating *p*-values and adjusting them for multiple comparisons is a fundamental aspect of many methods in statistical genomics. To mitigate inflated Type I error rates, *p*-values should be at least conservative. However, the statistical power may increase if these values are well-calibrated, balancing the control of false positives with the detection of true effects. The non-adjusted *p*-values computed by `cpam` are well-calibrated (i.e., within the uniformity envelope) for both short and long case-only time series (see Figures 5A and 5B), with similar results for case-control series (see Figure S1). For the pairwise models, we included both the complete set of *p*-values for all pairwise comparisons (pairwise-all), and a reduced set with only the minimum *p*-value per target (pairwise-min). Most methods are conservative (i.e., above the dashed line) in the range −log_10_ *p* ≥1.3 (*p* ≤ 0.05), but only `cpam`, tdeseq and pairwise-all are consistently well-calibrated across all simulations.

**Fig 5.**
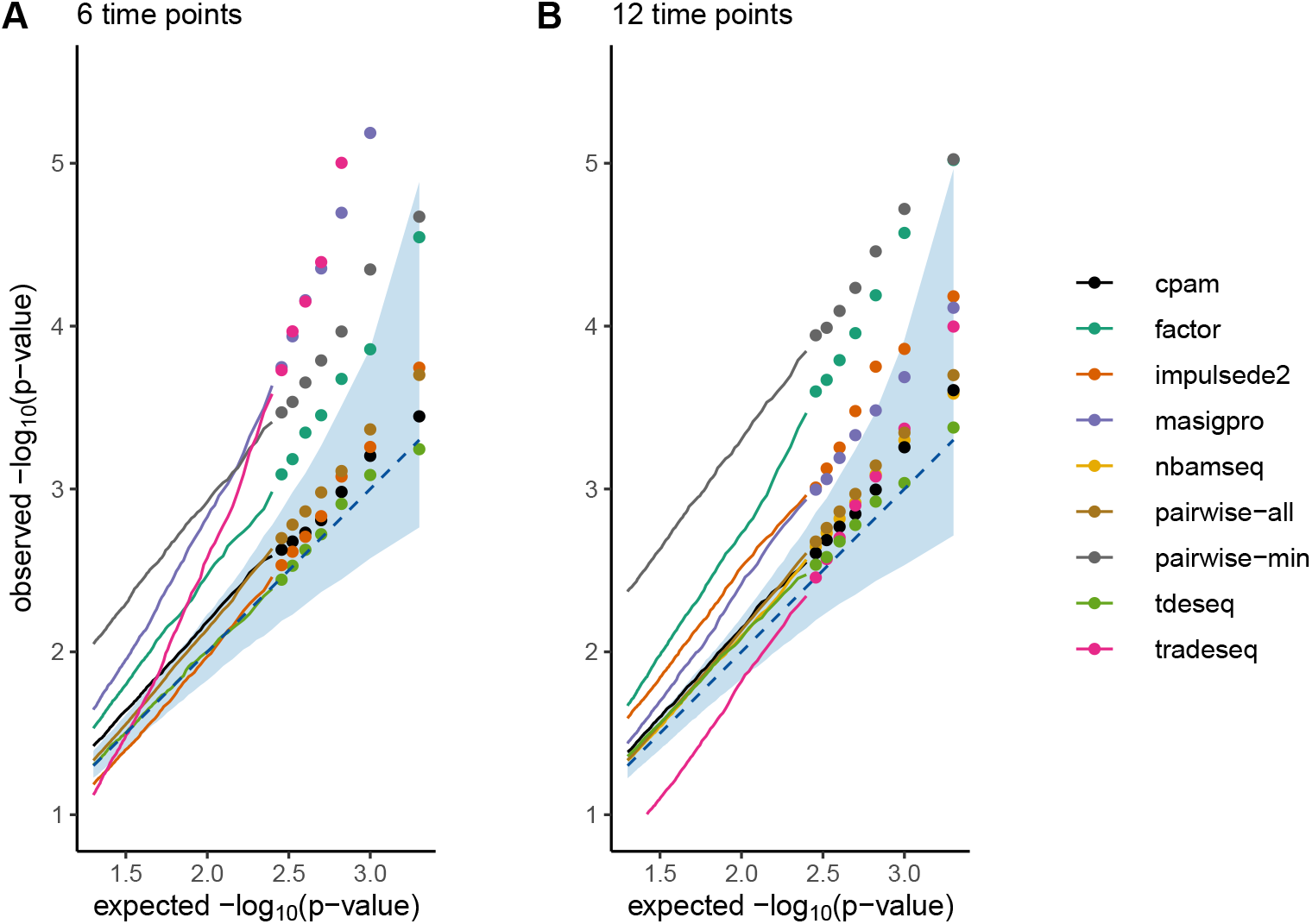
Quantile-quantile plots for time series methods comparing observed versus expected − log_10_(*p*-values) from null simulations for experiments with 6 (**A**) and 12 (**B**) time points. The diagonal dashed line represents perfect calibration where observed *p*-values match their theoretical distribution under the null hypothesis. The envelope shows the 95% confidence region expected under uniformity. `cpam` demonstrates well-calibrated *p*-values across both experimental designs, staying within the expected envelope, while several existing methods consistently deviate from the expected distribution.

### Detection of temporal differential gene expression

Statistical models for detecting differential gene expression should have high true positive rates (TPR) and low false discovery rates (FDR) at a set nominal FDR threshold. TPR is the proportion of true positives identified, while FDR is the proportion of false positives among discoveries. Figure 6 displays the TPR-FDR trade-off curves for increasing nominal FDR values, using simulated long series data with 10% true positives and 90% true negatives with maximal LFC = 1. See Figure S2 for short time series. For case-only (Figure 6A) and case-control (Figure 6B) time series, `cpam` out-performs the other methods by maintaining a higher TPR across relevant FDR levels. There are fewer curves plotted in Figure 6B as only a subset of the methods are applicable to case-control data. For case-only, only `cpam`, tdeseq, nbamseq, and trade-seq adequately control the FDR (i.e., the filled circles are close to the vertical dashed lines), and for case-control, only `cpam`.

**Fig 6.**
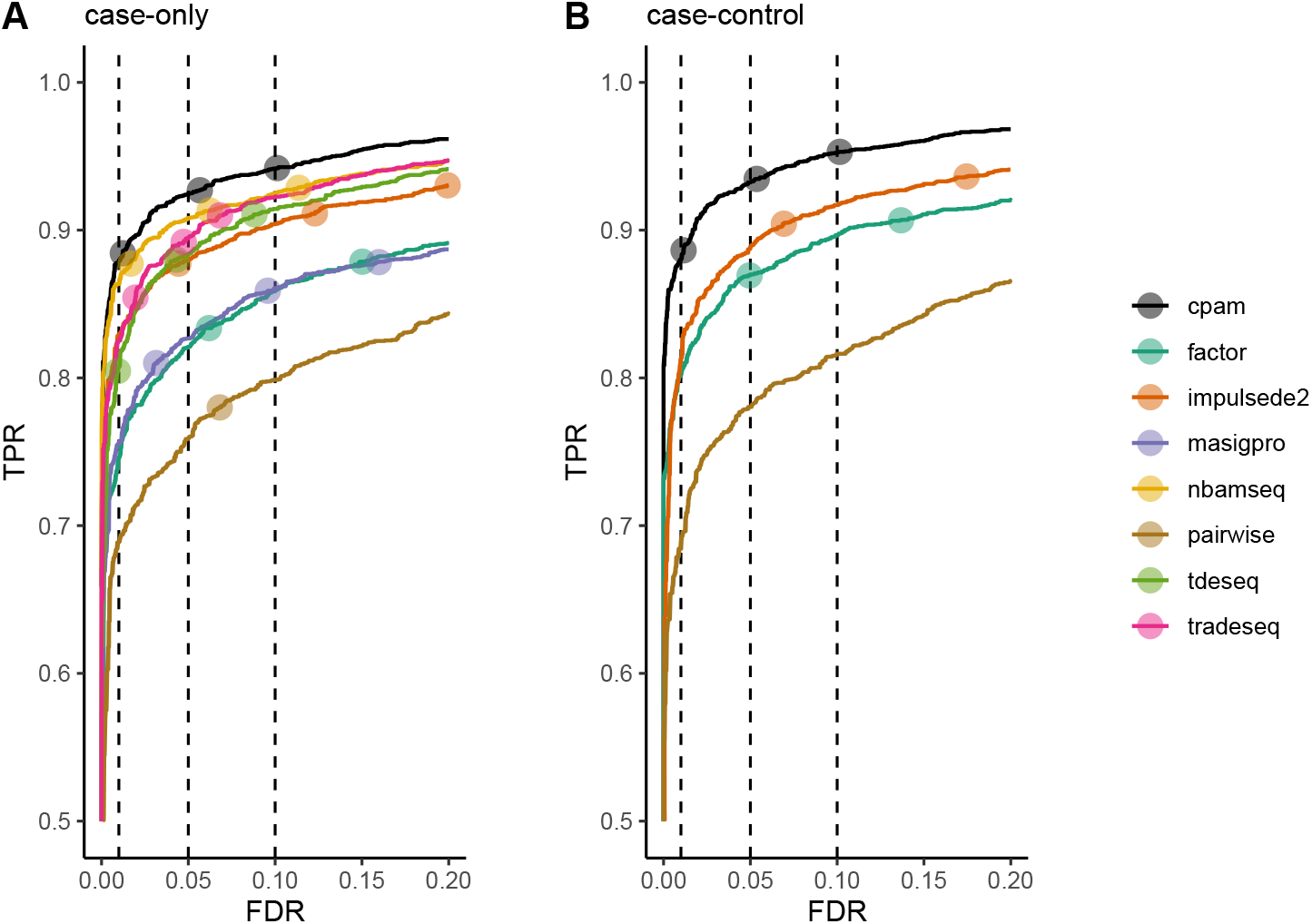
True positive rate (TPR) versus false discovery rate (FDR) for time series methods. The curves are computed from simulated data comprising 12 timepoints of three replicates for 3 *×* 10^4^ targets for case-only (**A**) and case-control (**B**) series. 10% of the targets are true positives. The three filled circles on each curve denote the actual TPR and FDR values that correspond to the nominal FDR thresholds 0.01, 0.05, and 0.1, ordered from left to right. Ideally, the three circles align with the corresponding reference lines of the observed FDR (vertical dashed lines). Curves that are closer to the top left corner have superior performance, finding more true positives while maintaining a low rate of false positives.

### Changepoint detection

To evaluate `cpam`’s changepoint detection, we simulated datasets with varying logfold changes (LFC) after the changepoint. We generated 30,000 targets across six timepoints with three replicates, comprising 10% non-null targets with changepoints. Non-null targets followed a pattern of (0, 0, 1, 2, 3, 4) (constant, then linear increase), scaled by LFC values ranging from 0.50 to 2.00 in increments of 0.25. This scaling determined the log_2_-fold increase between adjacent timepoints after the changepoint.

We evaluated changepoint detection methods within a complete workflow, including low-count filtering, dispersion estimation, and for `cpam`, restricting changepoint analysis to targets meeting an *α* = 0.05 FDR threshold. Performance was assessed by computing the bias (estimated minus true changepoint) and the proportion of correctly estimated changepoints for each LFC value. For bias calculations, the time between adjacent timepoints was set to one. For `cpam`, we tested both the one-standard error rule (1se, default) and the minimum scoring (min) methods for changepoint detection, see *Methods*. We also tested the pairwise method, where the changepoint was assigned to the earliest timepoint showing significance (*α* = 0.05) among all pairwise comparisons.

The results of the simulation study are shown in Figure 7. At small log-fold changes (LFC *<* 1), where early transcriptional changes typically occur, both `cpam` methods achieve substantially lower bias and higher accuracy than the pairwise approach. While the pairwise method improves with larger LFC, it never matches the performance of the other methods. For `cpam`, the spline-based modelling of the temporal autocorrelation accurately detects emerging changes in expression. This sensitivity to early molecular events is important for understanding causal mechanisms in gene regulatory networks. Except for LFC = 0.5, the one-standard error rule outperforms the minimum scoring method, indicating that the default method which aims to protect against overfitting leads to improved inferences.

**Fig 7.**
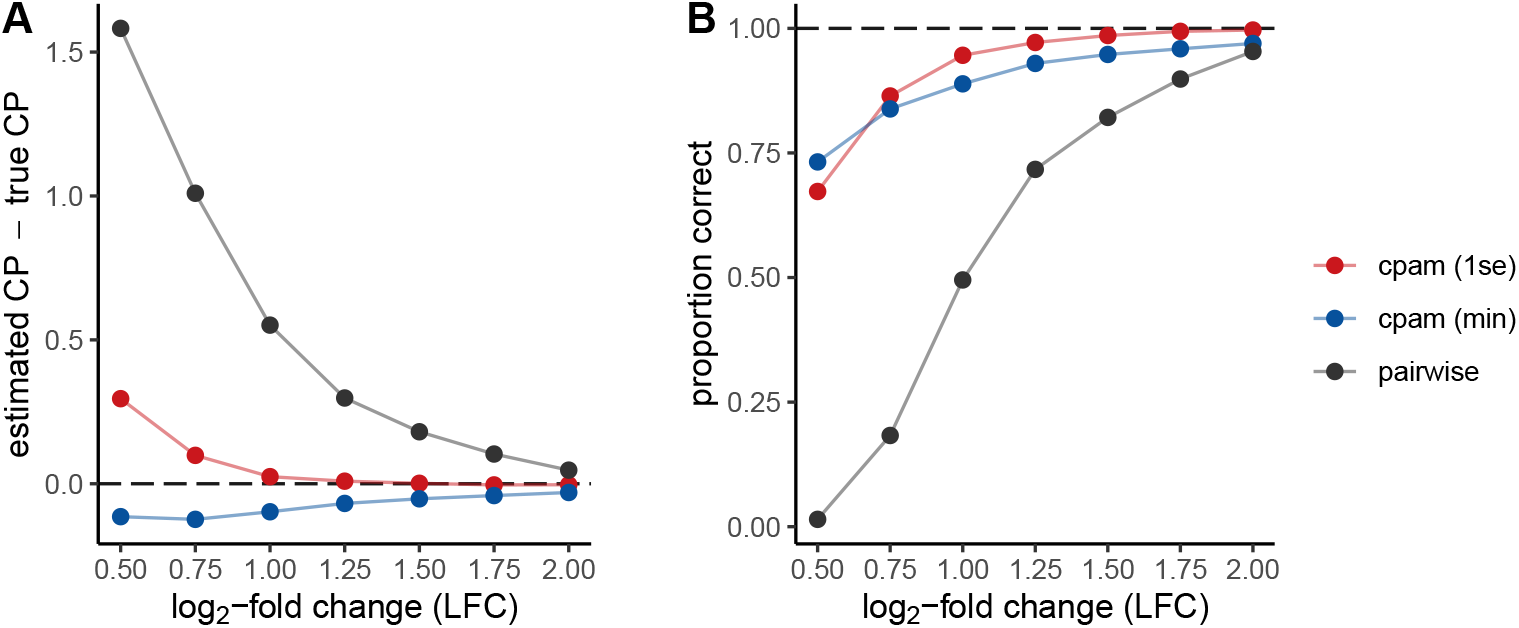
Evaluation of changepoint detection accuracy across different log_2_-fold changes (LFC). Data were simulated for 30 000 targets with 10% containing true changepoints, followed by linear increases scaled by different log_2_-fold changes (LFC).**A** Bias in changepoint estimation (estimated minus true changepoint), where values closer to zero indicate more accurate detection of the true changepoint timing. **B** Proportion of correctly estimated changepoints demonstrates method accuracy at each effect size.

### Novel features in human and Arabidopsis time course data revealed by application of cpam

We applied `cpam` to two publicly available data sets which have previously been analysed using alternative methods, to demonstrate the novel statistical and biological insights that `cpam` provides. In the first example, we analysed RNA-seq data from a time series of human embryo development from zygote through 2, 4 and 8 cell, morula and blastocyst stages (Torre et al., 2023). The embryonic genome activation (EGA) transition occurs after the 4-cell stage (t = 2.0 days). These data are ideally suited to highlight the powerful clustering capabilities of `cpam` from which we selected transcripts with an EGA change point (t = 2.0 days) and increasing concave shape. Crucially, the shape-constrained trends generated by `cpam` are easily sub-clustered based on user-defined factors such as rate of change and time to steady state (Figure 8A). Next, we illustrate application to one specific gene namely *PTRH2* (ENSG00000141378), a peptidyl tRNA hydrolase to show resolution at RNA isoform level. This gene has eleven isoforms, six of which passed the initial filtering stages, and `cpam` can model the expression shape for each isoform (Figure 8B). Importantly, different isoforms may belong to different expression clusters, showing the limitations of gene-level clustering and the importance of isoform-level clustering.

**Fig 8.**
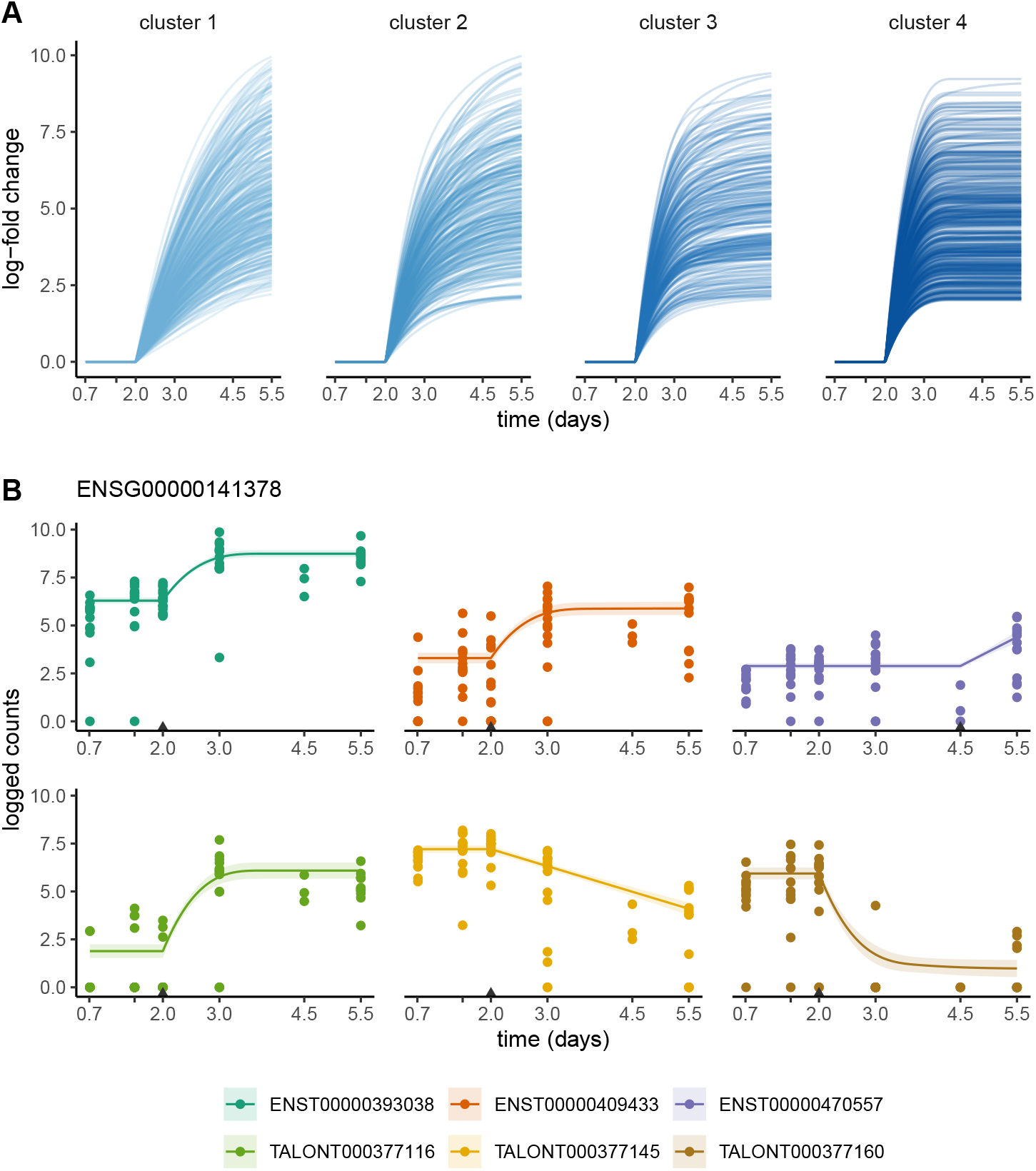
Application of `cpam` to RNA-seq time series of human embryo development.**A** RNA isoforms all with EGA changepoints (t = 2.0 days) and increasing concave shapes clustered by the ratio of log-fold changes between *t* = 3.0 and *t* = 5.5 h. **B** Shape-constrained models of RNA isoforms from the *PTRH2* (ENSG00000141378) gene

In the second example, we applied `cpam` to RNA-Seq data from an experiment in which Arabidopsis plants were transferred from low to very high (excess) light for 60 min (sampled at 0, 30 and 60 min) and returned to low light for 60 min (sampled at 67.5, 70, 90 and 120 min). Raw data were obtained from NCBI (Bioproject accession number: PRJNA391262) and trimmed based on quality using trimmomatic (Bolger et al., 2014). Kallisto (Bray et al., 2016) was used to quantify RNA isoform counts using 100 bootstrap replicates to estimate quantification uncertainty.

Application of `cpam` identified 11,594 differentially expressed transcripts (from 10,117 genes) compared to 13,572 gene-level DEGs previously identified (Crisp et al., 2017). The trend-based approach of `cpam`, which maintains a high true positive rate while accurately controlling the false discovery rate (Figure 6), finds fewer putative DEGs overall, but here it identified 910 additional ones. Among these new DEGs were two transcription factor genes. One is a *WRKY* gene (AT1G64140) showing transient upregulation (Figure 9A) as sought in the original analysis, while the other is *bZIP11* (AT4G34590) which shows sustained upregulation (Figure 9B) and has been previously characterised as responsive to energy status (Ma et al., 2011). These two genes produce single transcripts, but we found other genes which demonstrate transcript switching, involving alternative RNA splicing to produce different protein products (Figure 9C, 9D). The example in Figure 9D encodes a ubiqutin ligase *XBAT35* (AT1G26810) (Carvalho et al., 2012) in which only one isoform has a nuclear localisation signal. These examples demonstrate that `cpam` can find potentially important new genes and gene functions.

**Fig 9.**
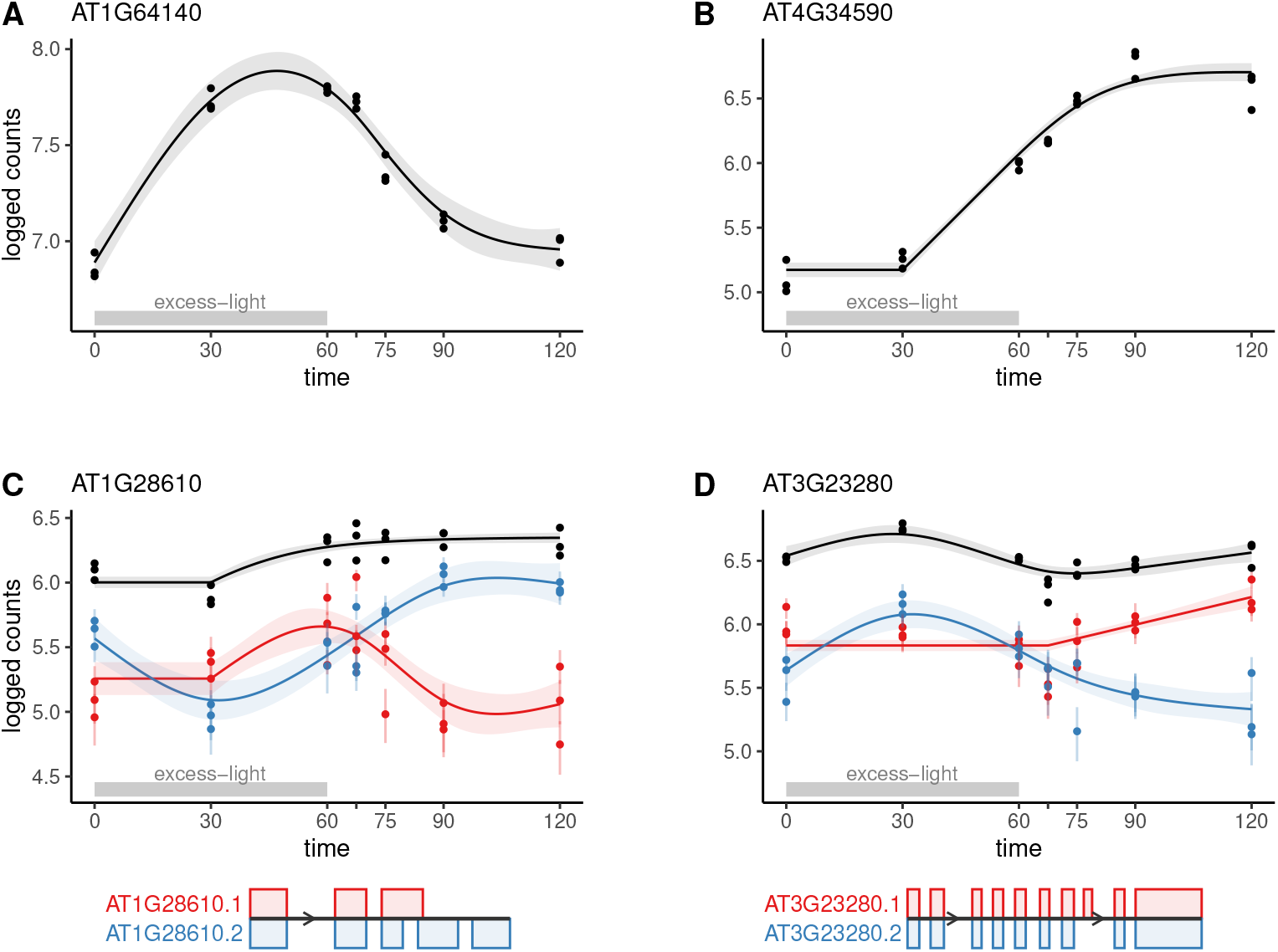
Application of `cpam` to RNA-seq time series of *Arabidopsis* plants treated with excess-light. Plants were treated with excess light for 60 min then returned to low light.**A** Expression profile of a *WRKY* gene (AT1G64140). **B** Expression profile of *bZIP11* (AT4G34590). **C** Expression of a lipase gene (AT1G28610) showing the gene-level profile (black) and profiles of two isoforms (red and blue). **D** Expression of ubiqutin ligase *XBAT35* (AT1G26810) showing gene-level profile (black) and two isoforms (red and blue). Gene structure schematics beneath panels C and D show the translated exon structures of the two isoforms in each case, illustrating that RNA isoforms produce different protein isoforms for each gene.

**Fig 10.**
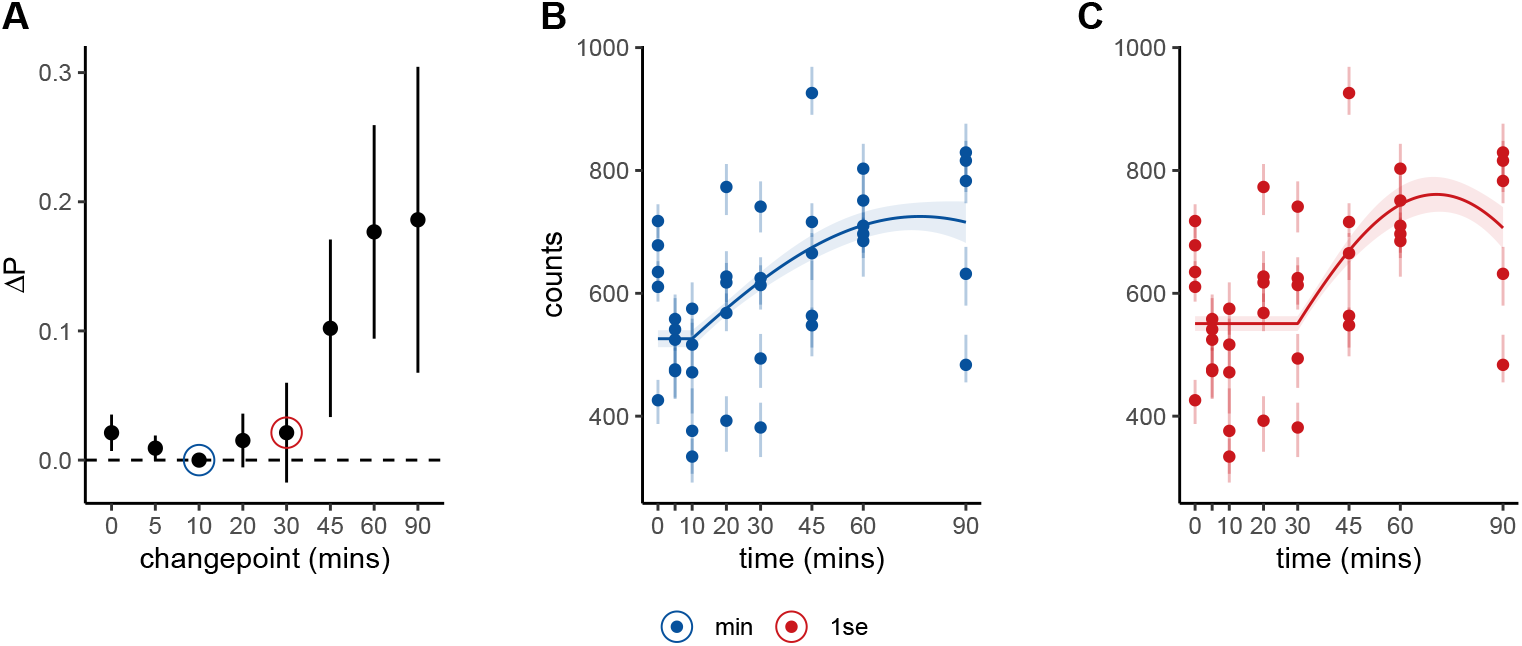
Illustration of changepoint model selection using the one-standard error rule.**A** Raw count data over time with fitted trajectory using the minimum scoring model (blue line) which places the changepoint at time 10. **B** The same data fitted using the one-standard error rule (red line), which selects a more conservative changepoint at time 30. **C** Differences in predictive criterion (Δ*P*) relative to the best-scoring model for each candidate changepoint, with vertical lines showing ±1 standard error. The one-standard error rule selects the latest changepoint (most parsimonious model) with a score within one standard error of the minimum (indicated by the horizontal dashed line), helping to protect against overfitting due to model-selection uncertainty.

## 3 Discussion

As large-scale omics experiments become increasingly accessible due to decreasing costs, there is a clear shift towards time-series experiments. Such experiments provide insights into critical biological processes including stem cell differentiation, embryogenesis, infection dynamics, ageing mechanisms, and responses to abiotic factors. These experiments span biological scales, from single cells to complex organisms and ecosystems, with implications for health care, agriculture, and environmental management. Here we show that `cpam` enables comprehensive temporal analysis by combining smooth temporal trends, transcript-and gene-level inferences accounting for quantification uncertainty, and biologically-motivated shape classification and changepoint estimation for clustering and pathway inference. This unified approach provides greater statistical power than existing methods while maintaining false discovery control, and enables detailed characterisation of temporal omics programs.

From statistical principles, trend-based modelling approaches should outperform pairwise comparisons by sharing information across time points, reducing multiple testing burden, and better reflecting the continuous nature of gene expression trajectories. Our results (Figure 6) confirm this expectation, in contrast to Spies et al. (2019) who found pairwise comparisons to be one of the best performing approaches for time series with up to 8 time points. This discrepancy may be explained by differences in simulated trends, where Spies et al. (2019) appear to have used temporal patterns that include large log-fold changes (LFC *>* 7) that can benefit pairwise detection (e.g., see Figure 7). Our simulations controlled the maximum LFC for all targets within a given simulation to assess model performance as a function of LFC.

Time series experiments aim to identify sequential gene expression patterns, triggered by environmental or developmental signals. The changepoint estimation of `cpam` identifies early gene expression changes with high confidence, even when the initial change is small in magnitude. This enables us to order change events in a chronological sequence, which in turn provides information about possible cause-and-effect relationships, and to infer pathways of responses.

The shape selection component of `cpam` provides a biologically interpretable framework to understand the dynamics of gene expression. By fitting predefined smoothed shapes that mirror common biological responses (e.g., sustained or transient changes), we create a standardised language for describing and comparing expression patterns.

Combined with changepoint detection, this approach reveals not only the timing of changes but also captures the rate and persistence of responses (e.g., distinguishing between rapid versus gradual changes, or temporary versus sustained responses). Although previous methods have introduced shape-based analysis (Fischer et al., 2018; Bar et al., 2022; Fan et al., 2024), our new method expands on these by incorporating the estimation of change points, target-specific smoothing, and transcript-level inferences that incorporate quantification uncertainty and permit the aggregation of *p*-values to the gene level for improved inferential power (Soneson et al., 2015).

The clustering approach of `cpam` can capture temporal dynamics of gene expression, by considering both the timing and pattern of gene expression changes. While two genes may share the same changepoint, they can exhibit different dynamics (e.g., a gene target of a transient transcription factor may show a slower rate of change; Figure 8). The approach in `cpam` is based on clustering genes by the modelled trends rather than by the expression counts themselves. This is particularly important as clustering purely on counts can allow spurious patterns to dominate the apparent dynamics of clustered expression patterns – even when they do not show statistically significant changes through time. The flexibility to choose the parameters by which gene expression clusters are defined enables `cpam` to create an almost limitless number of sub-clusters within each primary shape category. At the same time, `cpam` provides sensible clustering defaults (changepoint and shape) while allowing for user-defined customisation to explore more nuanced expression patterns.

An important application in modern omics programmes is the analysis of single-cell RNA sequencing (scRNA-seq) data. In this context, cells are first clustered into cell types based on their expression profiles, using methods such as PCA or UMAP (Hou et al.), and then assigned to inferred trajectories corresponding to different biological processes. Each cell is assigned a “pseudo-time” value, representing its progress along its trajectory (Saelens et al.). As individual cell data are typically noisy and sparse, cells on a given trajectory can be grouped into pseudo-time intervals and their expression counts aggregated, to create more robust “pseudobulked” measurements. These pseudobulked counts can then be analysed by `cpam` to characterise the temporal dynamics of gene expression along each trajectory, providing insight into cellular differentiation and developmental processes.

It is becoming increasingly apparent that a large proportion of eukaryotic genes produce two or more transcripts, known as RNA isoforms. Analysing gene expression at the isoform level is crucial because RNA isoforms produced from a single gene often exhibit different expression patterns across biological conditions and can differ in their stability, translatability, and encoded protein products. We have shown that `cpam` can readily identify significant genes in cases of isoform switching (Figure 9C, 9D). It is particularly important to undertake clustering analysis at isoform level because clustering at the gene level may be misleading for the thousands of genes with more than one transcript.

For time series experiments with a fixed sample budget, we recommend prioritising temporal resolution over replication. The smoothing splines in `cpam` leverage temporal autocorrelation to share information across time points, enabling accurate inference even with fewer replicates. While we suggest a minimum of 5 time points to access the advantages of splines over pairwise comparisons, the optimal number of replicates depends on sampling frequency—closely spaced measurements may need only 1-3 replicates, while widely spaced measurements may require 3 or more. The definition of “close” versus “wide” spacing ultimately depends on the rate of change in the biological process under study. An interesting investigation for the future may be to determine optimal sampling strategies for different types of temporal dynamics.

We envisage some extensions of the `cpam` method and software to be considered in future work. One interesting avenue has emerged from our investigations of changepoint additive models for RNA-seq data, which revealed that RNA isoforms often exhibit differential variance post-changepoint. For example, certain isoforms become more tightly regulated (i.e., less variable) post-changepoint despite an increase in their mean expression. This is in keeping with observations that changes in variance can be used to identify cancer-related genes in humans that are missed by standard differential expression analysis (Roberts et al., 2022). The present work could be extended to allow target-specific models of the variance (dispersion) parameter using the framework of generalised additive models for location, scale, and shape (GAMLSS, Stasinopoulos and Rigby, 2007; Wood, 2017). Additionally, a current limitation of `cpam` is the restriction to a single changepoint. We imposed this constraint largely for computational reasons, but an additional changepoint may be helpful, for example, to identify when a target attains a stabilised expression level.

Our new package `cpam` provides a comprehensive framework for analysing time series omics data that combines statistical rigour with practical utility. The method leverages modern statistical approaches while remaining user-friendly, through sensible defaults and an interactive interface. Researchers can directly address key questions in time series analysis—when changes occur, what patterns they follow, and how responses are related. While we have focused on transcriptomics, the framework is applicable to other high-dimensional time series measurements, making it a valuable addition to the omics analysis toolkit.

## 4 Methods

### Filtering low-count targets

To increase statistical power, low-count targets are filtered from the data prior to the estimation of normalisation factors and dispersions. The default filtering function retains a target if at least 60% (min prop = 0.6) of the samples have a minimum of 5 (min reads = 5) reads at one or more timepoints. Users can manually set min prop and min reads.

### Dispersion estimation

Dispersions are estimated using the approximate empirical Bayes method implemented in the R package edgeR (McCarthy et al., 2012). The method fits a trend line to the relationship between the mean expression levels and a corresponding initial set of independent dispersion estimates. In `cpam`, the design matrix for these initial estimates is constructed from orthogonal quadratic polynomials in the time variable. The final dispersion estimate for each target is obtained by shifting, or regularising, the initial estimate towards the trend line, where the amount of regularisation depends on the number of samples, the mean count, the consistency of the target-wise dispersions, and distance of the initial estimate from the trend line. The use of empirical Bayes regularisation can be optionally switched off.

### Quantification uncertainty

Unlike gene counts, the quantification of isoform reads can be highly uncertain because of ambiguities in mapping reads to specific isoforms. To estimate the quantification uncertainty, computationally efficient pseudo-alignment tools such as *kallisto* (Bray et al., 2016) and *salmon* (Patro et al., 2017) allow for the rapid simulation of boot-strapped technical replicates. Although the use of these tools to generate isoform counts is common, the bootstrap replicates themselves are not often used to inform downstream analyses, perhaps due to limited software options. Here we apply the method recently developed by Baldoni et al. (2023) to estimate a bootstrapped overdispersion coefficient 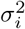 for each target *i*. The coefficient is used to rescale the counts, or ‘divide out’ the quantification uncertainty, before model fitting. We refer the reader to the original paper for details of the method.

When the data supplied are not a fixed count matrix but outputs from external quantification software, `cpam` imports them using the R package *tximport* (Soneson et al., 2015) and optionally applies the overdispersion scaling method if bootstrap replicates are present. When applied, the scaled counts are used for all model fitting stages including the estimation of *p*-values, changepoints and shapes; however, the default plotting functions show the predicted trends and the observed data without the scaling.

### Model fitting

The temporal smoothing splines introduced in Equations (3) and (4) are constructed using a set of *k* basis functions, *f*_*n*_, *n* = 1, 2, .., *k*, for which a single spline can be expressed as

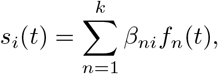

where *i* indexes the target and *β*_*ni*_ are the parameters to be estimated. For the regular smoothing splines used to estimate *p*-values and changepoints in `cpam`, the basis functions are thin plate splines and *k* is set to the number of timepoints in the series Wood et al. (2016). For shape-constrained models, the basis functions are derived from *P* -splines, and the coefficients *β*_*ni*_ are subject to shape-specific sign and ordering constraints which are imposed via re-parametrisation (Pya and Wood, 2014). For all models, the parameters ***β***_*i*_ = (*β*_*ni*_) are estimated using penalised regression, where the strength of the penalty depends on the curvature of the basis functions and an estimated smoothing parameter *λ*_*i*_ ≥ 0. The model can be understood from a mixedeffects or Bayesian perspective, where the penalisation is expressed through the prior distribution,

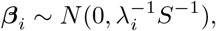

with the elements of the penalty matrix 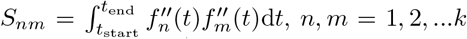, derived from the second derivatives of the basis functions. All targets use the same set of basis functions, which depend only on the observed timepoints, and therefore share a common *S*. The smoothing spline models in `cpam` are fit with the R packages mgcv (Wood et al., 2016) and scam (Pya, 2024) using either restricted maximum likelihood (REML) or generalised cross validation (GCV) to estimate *λ*_*i*_.

### Model selection

Model selection for the estimation of changepoints and shapes makes use of the pointwise estimates of a predictive criterion *P* , where *P* is either GCV or AIC. For a set of models indexed by *k* = 1, …, *K*, the criterion estimate is written as

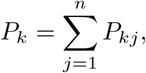

where *n* is the number of samples and *P*_*kj*_ is the predictive log-likelihood of the *j*th sample for model *k*. Both criteria are derived from a single model fit by computing the pointwise log-likelihood of the data and applying a correction based on the effective degrees of freedom (EDF) of the model, as calculated by mgcv. For AIC, the correction is additive, while for GCV it is multiplicative (Wood, 2017). An advantage of computing the criterion through pointwise estimates is that model comparisons based on differences Δ*P*_*k*_ = *P*_*k*_ − min_*k*_ (*P*_*k*_) can be accompanied by estimates of the standard error 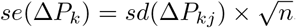.

Figure 10A shows a plot of Δ*P*_*k*_ estimates and their standard errors, where *k* denotes the changepoint characterising each candidate model. For this target, the model with a changepoint at 10 has the best score (Figure 10B), but the models with changepoints at 20 and 30 (Figure 10C) have comparable scores when taking into account the uncertainty of the score differences (i.e., the standard errors of the differences overlap zero). To mitigate overfitting due to model-selection uncertainty, `cpam` applies the modified one-standard error rule (Yates et al., 2021) and chooses the simplest (smallest EDF) model within 1*se* of the best scoring model.

For the changepoint models, the model with the latest changepoint will usually have the smallest EDF, where a changepoint at the final timepoint is simply the null model. For selection among shapes, convex and concave are the most complex, followed by their monotonic versions, and then linear trends. Within this nested hierarchy, model complexity is ranked by EDF. Instead of the one-standard error rule (default), users can opt to use the minimum scoring model.

## Acknowledgements

This research was funded by the Australian Research Council, Grant CE200100015. We are grateful for the use of publicly available datasets for allowing the exposition of the capabilities of `cpam`.

## Supplementary Material A

**Fig S1.**
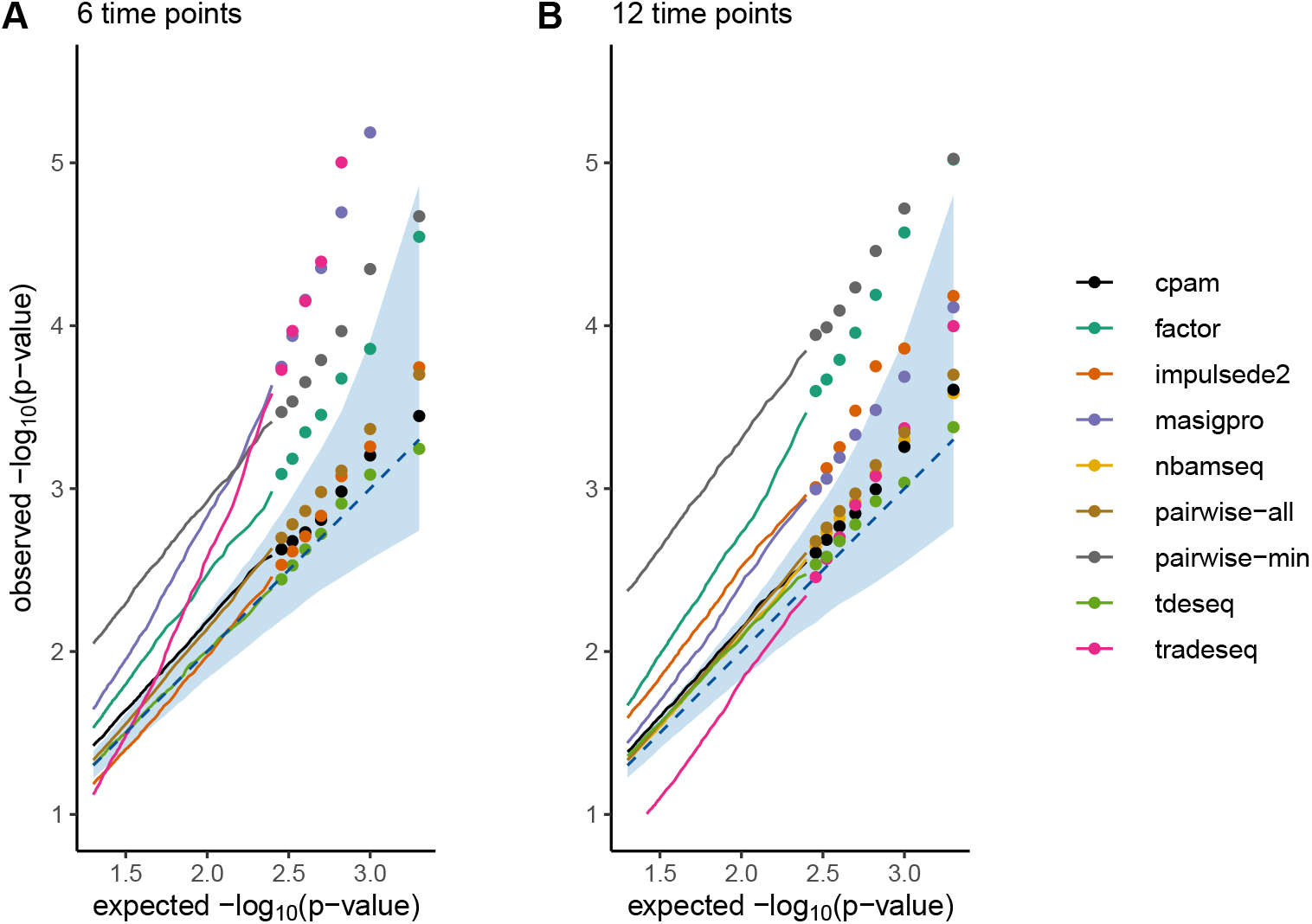
Distribution of *p*-values under the null hypothesis. Case-control.

**Fig S2.**
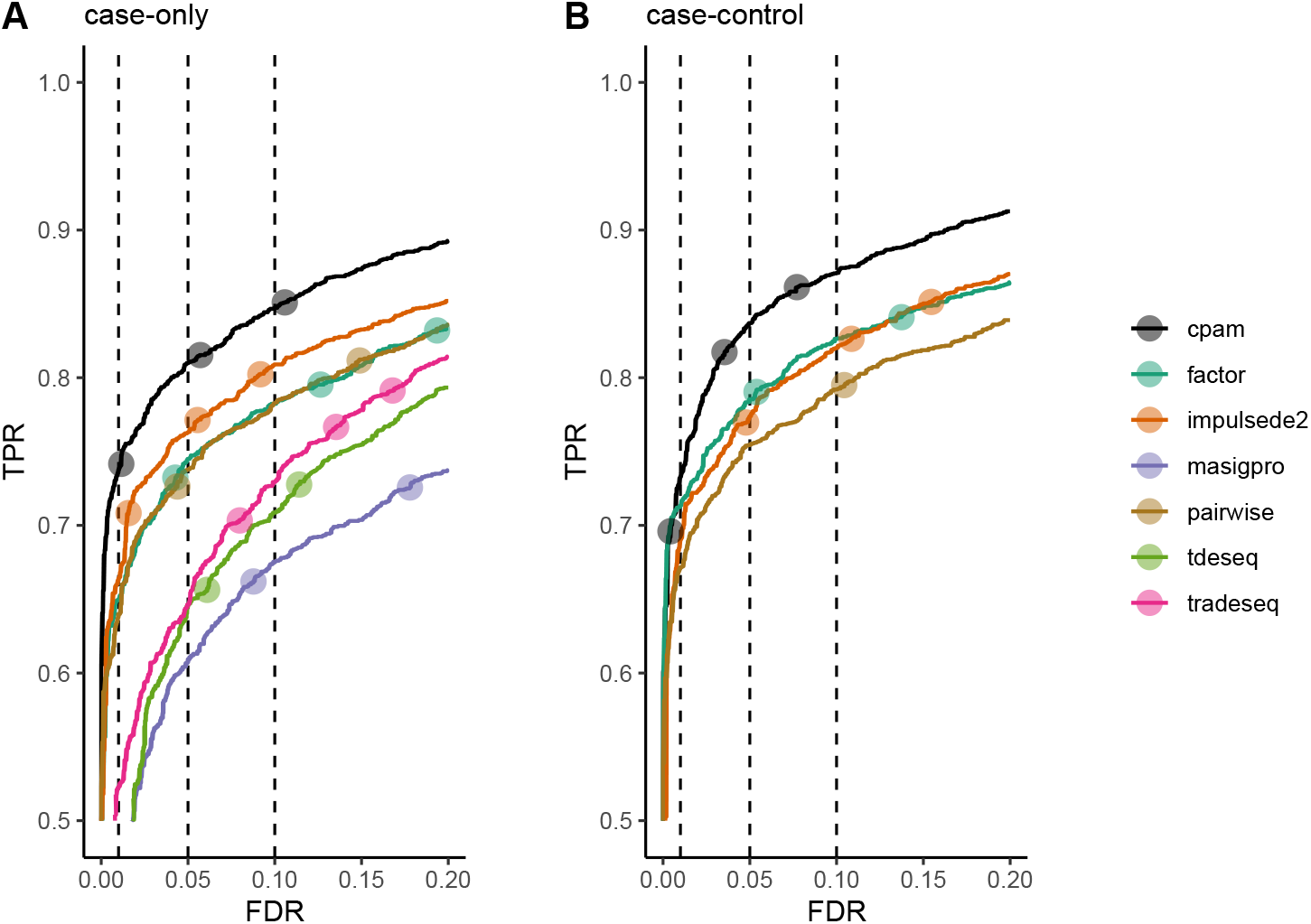
True positive rate (TPR) versus false discovery rate (FDR) for time series methods. The curves are computed from simulated data comprising 6 timepoints of three replicates for 3 × 10^4^ targets. 10% of the targets are true positives. The three filled circles on each curve denote the actual TPR and FDR values that correspond to the nominal FDR thresholds 0.01, 0.05, and 0.1, ordered from left to right. Ideally, the three circles align with the corresponding reference lines of the observed FDR (vertical dashed lines). Curves that are closer to the top left corner have superior performance, finding more true positives while maintaining a low rate of false positives.

## Notes

### Competing Interest Statement

The authors have declared no competing interest.

